# Is Boldness in Climbing Perch (*Anabas testudineus*) Sensitive to the Apparatus Used for Measurement?

**DOI:** 10.1101/2020.07.06.189092

**Authors:** Prantik Das, V V Binoy

**Affiliations:** National Institute of Advanced Studies (NIAS), Indian Institute of Science (IISc) Campus, Bangalore 560012, India

**Keywords:** Exploratory behaviour, neophobia, boldness, open field, repeated exposure, measuring behaviour, scotophobia, swimway

## Abstract

Swimway and open field are the two popular apparatus used for measuring boldness - the propensity to take risky decisions - in various piscine species. The present study compared boldness exhibited by an air breathing freshwater fish climbing perch in a swimway, rectangular open field, intermediate stages between these two apparatus and circular open field. Impact of the modification of the start chamber by providing substratum made up of cobbles and covering the water surface using water plant hydrilla, alone and in combinations on the boldness was also tested. Our results revealed that the apparatus has a significant impact on the boldness in climbing perch. The presence of a shelter in the experimental arena (swimway) and cobble substratum in the start chamber of the apparatus were found to be reducing boldness in this fish, while hydrilla cover on the water surface neutralised the impact of cobble substratum. Repeated exposure and resultant familiarity with the arena increased boldness of climbing perch but the pattern of modification of this behavioural trait exhibited during the course of experiment was divergent across the instruments. These results point towards the need for critically analyzing the influence of instruments used for measuring various behavioural traits and considering biological needs of the subject species while designing the apparatus.

## Introduction

The boldness - propensity to take risky decisions - and its behavioural and physiological correlates are being studied extensively in various fish species due to the close association between this trait and the fitness (White et al., 2013; Conrad et al., 2011; Pearish et al., 2019). The bold individuals with an enhanced tendency to take risk decorate the dominant positions and influence the dynamics of a shoal, establish new colonies and exhibit enhanced cognitive capacities (Trompf and Brown, 2014, Cote et al. 2011). However their reduced fear could put them in danger also; bold individuals may end up as easy prey and may even perish by eating unsafe and unfamiliar materials in novel habitats (Sundström and Johnsson, 2001). In fish, trait boldness is measured as the latency to initiate exploration of a novel area or an object (Conrad et al., 2011; Toms, 2010). In novel situations bold individuals take control over the fear (neophobia) and regain normal rhythm of activities quickly. Meanwhile in a similar situation their shy counterparts exhibit higher levels of freezing and startling responses and take a longer duration to overcome neophobia (Budaev et al., 1999). Although a wide range of methodologies and apparatus are being utilized for quantifying boldness in fishes; ‘open field*’* and swimway (emergence test) are the trendiest (Warren and Callaghan, 1976; Polverino et al., 2016; Toms et al., 2010; Yoshida et al., 2005; Brown et al., 2005).

The open field consists of a well illuminated homogeneous open area, either circular or rectangular in shape and varying in size according to the species being tested. Here, subject fish is introduced into the well-lit central area of the apparatus and the latency to initiate exploration is recorded. The bold individuals are seen to start exploration soon after their entry into the experimental arena, while shy ones exhibit prolonged latency to move around (Warren and Callaghan, 1976). However, many researchers believe that open field could force the subjects to move, induces hyperactivity and startling response and separating the effect of compulsion by environment from the actual inclination for exploration is very difficult when tested using this apparatus (Walsh and Cummins, 1976; Gould et al., 2009). According to Yoshida et al. (2005) swimway apparatus consisting of a shelter (opaque chamber) connected to an open area through a door is a better instrument for measuring boldness in fishes. Here the subject fish released into the start box, which acts as a shelter, has the freedom to stay inside or explore the open area outside which avoids the compulsion for initiating activity in a novel environment (Mazue et al., 2015). Akin to the open field, in this apparatus also bold individuals emerge from the start chamber quickly and extend their activities to the open area in comparison to the shy ones (Brown et al., 2007; Toms et al., 2010). The modified version of swimway is also being utilized for quantifying the boldness of various piscine species in their natural habitat (Brown et al., 2005). However Burns (2008) has a different opinion about the swimway apparatus; his study revealed that in this apparatus also the subject fish exhibits fear and startling response such as darting immediately after the gate was opened.

An exploration of the literature on the measurement of boldness in fishes would reveal that some species modified their tendency to explore novel areas in response to the alteration in the properties of the apparatus, while others did not. For instance, in the fry of brown trout (*Salmo trutta*) a minor modification like changing the size of the gate connecting shelter and open area of the swimway apparatus was found influencing the emergence latency (Näslund et al., 2015). Whereas boldness of the eastern mosquitofish, *Gambusia holbrooki*, remained independent of the spatial properties (size) of the apparatus utilised (Polverino et al., 2016). These results point to the need for quantifying and comparing boldness exhibited by a fish species in an open field, swimway and the instruments representing stages in between to get a complete picture of this vital behavioural trait. Unfortunately, although boldness is being studied in a large number of species only a few studies focusing on this important aspect have appeared in the literature till date (Näslund et al., 2015). The present study examined the influence of the instruments used for measuring boldness in an air breathing freshwater fish climbing perch. We tested the boldness of this species in the swimway, rectangular open field, two transitional stages between these apparatus and the circular open field. Scope of the current investigation also included understanding the influence of the modification of the start chamber of the swimway and repeated exposure for five consecutive days on the boldness of this ecologically and economically important species.

## Materials and methods

Climbing perch (*Anabas testudineus* Bloch) is a medium-sized, omnivorous, shoal-living fish inhabiting a wide variety of freshwater ecosystems in India and several south-east Asian countries (Talwar and Jhingran, 1991; Binoy and Prasanth, 2016). Being bestowed with labyrinthine organs for breathing atmospheric air, this fish could tolerate extreme environmental conditions and their populations have been reported from a wide variety of ecosystems including areas of saline intrusion and sewages (Binoy and Prasanth, 2016). This fish is a delicacy in many countries and its population is experiencing steady decline due to the over exploitation (Kohinoor et al., 2012). However in countries such as Australia this fish has emerged as an invasive species threatening local aquatic biodiversity (Hitchcock, 2008). Unfortunately, till date only a handful of studies have focused on the boldness in this species (Binoy and Thomas, 2003; Binoy, 2015; Binoy et al., 2019) though this information could be vital in developing strategies to conserve, culture and control this species.

### Focal species and husbandry

The subject populations of the climbing perch were collected from the irrigation canals of the Karalam (10.41°N; 76.19° E) region of the *Kole* paddy fields, Kerala State, India, with the support of expert fishermen. In the laboratory, fish of standard length 62.4 ± 26.0 mm (mean ± SD) were sorted and kept in groups of six in aquaria (45 cm × 22 cm × 22 cm). Water temperature was maintained at 25 ± 1°C and light hours at 12L:12D. Three sides of all aquaria were covered with black paper while steel grids placed on the top prevented the fish from jumping out. The fish were fed with commercial food pellets twice daily *ad lib*. and unused pellets were siphoned out after 30 min. The subject fish were given 7 days for acclimation with the laboratory conditions before testing.

### Experiment 1 Boldness and Apparatus Apparatus – 1: Swimway

The swimway apparatus (Yoshida et al., 2005) was an aquarium (60 × 32 × 32 cm) divided in two chambers, A (20 × 32 × 32 cm) and B (40 × 32 × 32 cm), using an opaque Plexiglass sheet with a guillotine door (8 × 4 cm) in the middle. The chamber A (which functioned as the start box) was covered from the top using an opaque acrylic sheet to provide shade for the subject fish staying inside. The water level was kept at a height of 28 cm. All the four sides of the apparatus were covered using black paper to avoid any sort of external interference.

### Apparatus – 1a and 1b

These apparatus were the intermediate stages between the swimway and open field. The swimway (apparatus 1) was modified as follows:

#### Apparatus 1a)

Here the opaque acrylic sheet covering the start box (chamber A) from the top was not present. This arrangement reduced the shade inside the start chamber hence its refuge value.

#### Apparatus 1b)

Here the opaque acrylic partition separating the chambers A and B in the *Apparatus 1a* was replaced with a transparent one with the gate.

### Apparatus 2: Rectangular open field

The apparatus 1a was converted into the rectangular open field by removing the partition wall separating the chambers A and B. Here the experimental arena was a homogenous area without any shade or refuge.

### Apparatus 3 Circular open field

A circular tank (inner radius 30 cm and height 35 cm) filled with water up to a height of 28 cm was used as the circular open field.

All these five apparatuses were illuminated with compact fluorescent lamps (20W) positioned at a height of 40 cm above the water surface.

### Measuring boldness (General Protocol)

#### Apparatus 1, 1a and 1b

Individual subject fish were introduced into chamber A and the guillotine door was opened after two minutes given for the acclimation. The latency to initiate the exploration of chamber B, defined as the time taken by the climbing perch to come out of chamber A through the door provided (Binoy, 2015), was recorded. The fish failed to come out of chamber A after 6 minutes were given a ceiling value of 360 seconds.

#### Apparatus 2 and 3

Subject climbing perch were introduced into the central area of the open field individually. Initiation of exploration of the fish in the open field, covering a distance of 3 body lengths, was taken as the measurement of boldness. Here also the individuals that continued to be stagnant in the experimental arena during the test time (6 minutes) were allocated a ceiling value of 360 seconds.

### Experiment 2: Boldness and repeated exposure

Influence of the repeated exposure to the experimental arena on boldness of climbing perch was studied using three different apparatus viz. swimway, open field (circular) and ‘apparatus 1b’. Here subject fish were tested individually once daily by following the protocols mentioned above for five days consecutively. We tested separate groups of fish in each apparatus.

### Experiment 3: Modified refuge and the boldness

This experiment explored the question, could the presence of floating vegetation and cobble substratum in the start box (chamber A) influence the boldness of climbing perch? Here we used apparatus 1b and subject fish were given experience with the experimental arena for five days as described in experiment 3, prior to the testing. Each of the 18 subject fish were tested individually in 3 different situations; start chamber with cobble (grain size range = 60-85 mm) substratum, water surface of the start chamber covered with a thick pack of water plant (*hydrilla verticillata*; in this experiment the substratum was glass) and a combination of cobble substratum and water plant. The subject fish was always introduced into chamber A and the boldness was measured following the general protocol. We took a lottery to decide the sequence of the exposure.

## Analysis

The data obtained from experiments 1 and 2 (for swimway and open field) was not following normal distribution even after transformation. Hence, we used nonparametric statistics Kruskal Wallis test (cross instrument comparison), Friedman’s repeated ANOVA (repeated exposure in experiment 2 for swimway and open field) and Dunn’s test (*post hoc* analysis). The parametric test repeated measures ANOVA followed by *post hoc* analysis (Tukey’s test) was used in cases where data was following normal distribution (repeated exposure in apparatus 1b and experiment 3).

## Result

### Experiment 1: Boldness and Apparatus

Our results revealed that the climbing perch exhibited a significant variation in the latency to initiate exploration of the novel area when tested using different apparatus (Kruskal Wallis test; *χ*^2^ = 34.91, p < 0.0001). Further analysis revealed that the boldness exhibited in the swimway was different from all other apparatus except circular open field (Dunn’s test; Table 1). Similarly the latency to start activity in the apparatus 1b was different from the rectangular and circular open fields (Fig. 1).

**Table 1.** Boldness exhibited by the climbing perch in different apparatus tested - *Post hoc* analysis (Dunn’s test). Statistical significance is denoted as * = p<0.05, ** = p<0.01, *** = p<0.001

|  | Apparatus 1a | Apparatus 1b | Open field<br>(Rectangular) | Open field<br>(Circular) |
| --- | --- | --- | --- | --- |
| **Swimway (With**<br>**shade on**<br>**chamber A)** | $\chi^2 = -4.35$<br>$p = 0.0001^{***}$ | $\chi^2 = -5.48$<br>$p = 0.0001^{***}$ | $\chi^2 = -2.86$<br>$p = 0.43^*$ | $\chi^2 = -2.42$<br>$p = 0.07$ |
| **Apparatus 1a** | — | $\chi^2 = 1.25$<br>$p = 1.00$ | $\chi^2 = -1.49$<br>$p = 0.67$ | $\chi^2 = -1.93$<br>$p = 0.26$ |
| **Apparatus 1b** | — | — | $\chi^2 = -2.70$<br>$p = 0.03^*$ | $\chi^2 = -3.13$<br>$p = 0.01^{**}$ |
| **Open field**<br>**(Rectangular)** | — | — | — | $\chi^2 = 0.44$<br>$p = 1.00$ |

**Figure 1.**
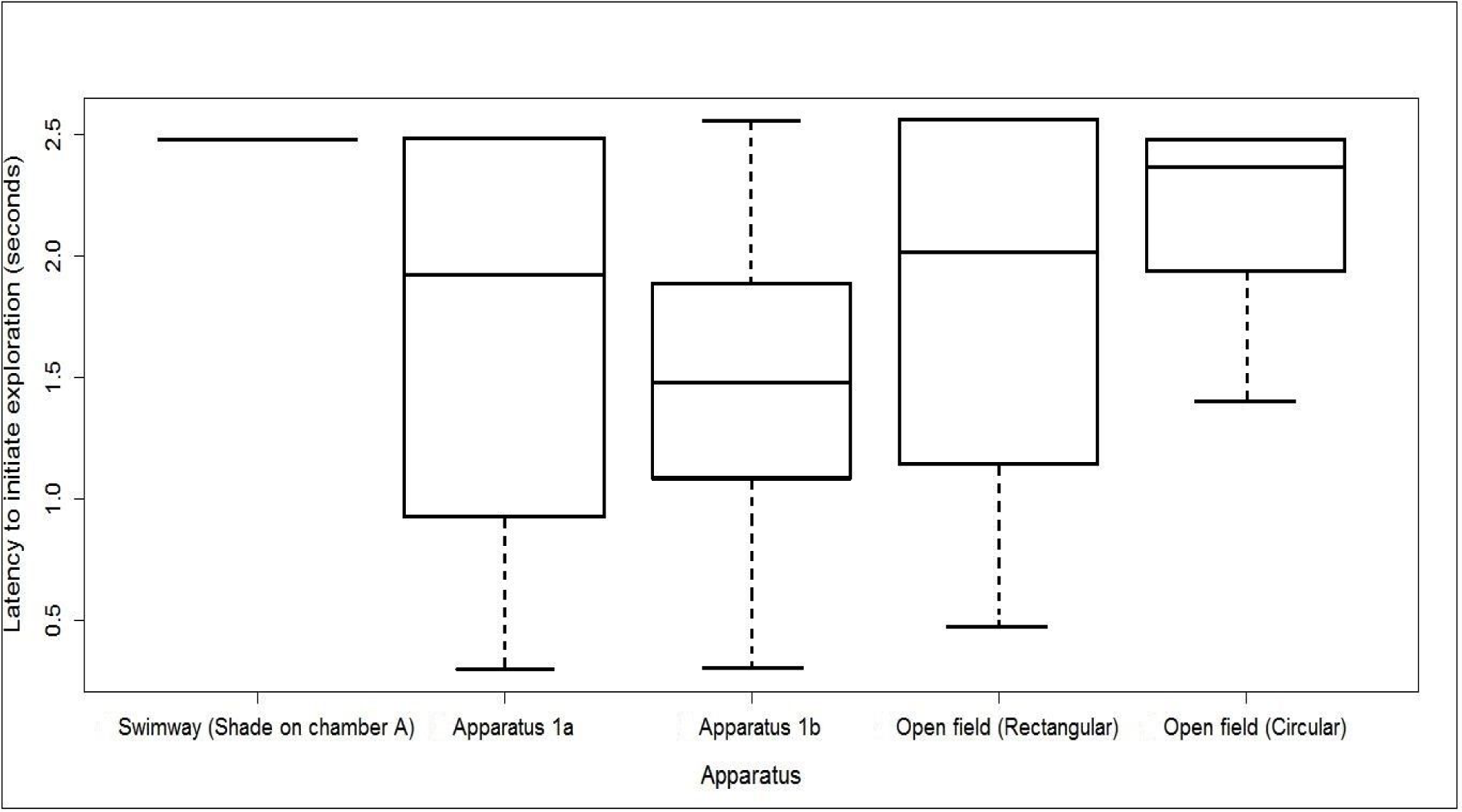
Boldness exhibited by the climbing perch in different apparatus tested. The data is log converted.

### Experiment 2: Boldness and repeated exposure

The repeated measurement of boldness using swimway (Friedman’s repeated ANOVA; *χ*^2^ = 11.44, p = 0.02; Fig. 2), and circular open field (Friedman’s repeated ANOVA; *χ*^2^ = 4.89, p = 0.29; Fig. 4) revealed a positive influence of the experience with the experimental arena on this behavioural trait. However the patterns of modification exhibited by the boldness in response to the repeated exposure for five consecutive days in these two apparatus were not similar. In the swimway, the latency to initiate exploration exhibited on the 1^st^ day was significantly different from the all other days, except 4^th^ day (Table 2), while in the circular open field such a variation was seen between day 4 and day 5 only (Table 3). Interestingly, in apparatus 1b no significant difference was noticed in the boldness exhibited by the climbing perch in any of the five days of repeated exposure (Repeated measures ANOVA; F_14_ = 1.07, p = 0.38; Fig. 3)

**Table 2.** Influence of the repeated exposure for five consecutive days on the boldness of climbing perch in swimway apparatus - *Post hoc* analysis (Dunn’s test). Statistical significance is denoted as * = p<0.05, ** = p<0.01, *** = p<0.001

|  | **Day 2** | **Day 3** | **Day 4** | **Day 5** |
| --- | --- | --- | --- | --- |
| **Day 1** | $\chi^2 = 2.77$<br>$p = 0.02^*$ | $\chi^2 = 2.86$<br>$p = 0.02^*$ | $\chi^2 = 2.12$<br>$p = 0.16$ | $\chi^2 = 2.92$<br>$p = 0.01^*$ |
| **Day 2** | — | $\chi^2 = 0.06$<br>$p = 1.00$ | $\chi^2 = -0.66$<br>$p = 1.00$ | $\chi^2 = 0.20$<br>$p = 1.00$ |
| **Day 3** | — | — | $\chi^2 = -0.72$<br>$p = 1.00$ | $\chi^2 = 0.16$<br>$p = 1.00$ |
| **Day 4** | — | — | — | $\chi^2 = 0.85$<br>$p = 1.00$ |

**Table 3.** Influence of the repeated exposure for five consecutive days on the boldness of climbing perch in circular open field - *Post hoc* analysis (Dunn’s test). Statistical significance is denoted as * = p<0.05, ** = p<0.01, *** = p<0.001

|  | **Day 2** | **Day 3** | **Day 4** | **Day 5** |
| --- | --- | --- | --- | --- |
| **Day 1** | $\chi^2 = 0.74$<br>$p = 1.00$ | $\chi^2 = 1.51$<br>$p = 0.65$ | $\chi^2 = 0.97$<br>$p = 1.00$ | $\chi^2 = 2.48$<br>$p = 0.04^*$ |
| **Day 2** | — | $\chi^2 = 0.76$<br>$p = 1.00$ | $\chi^2 = 0.22$<br>$p = 1.00$ | $\chi^2 = 1.73$<br>$p = 0.41$ |
| **Day 3** | — | — | $\chi^2 = -0.53$<br>$p = 1.00$ | $\chi^2 = 0.97$<br>$p = 1.00$ |
| **Day 4** | — | — | — | $\chi^2 = 1.51$<br>$p = 0.65$ |

**Figure 2.**
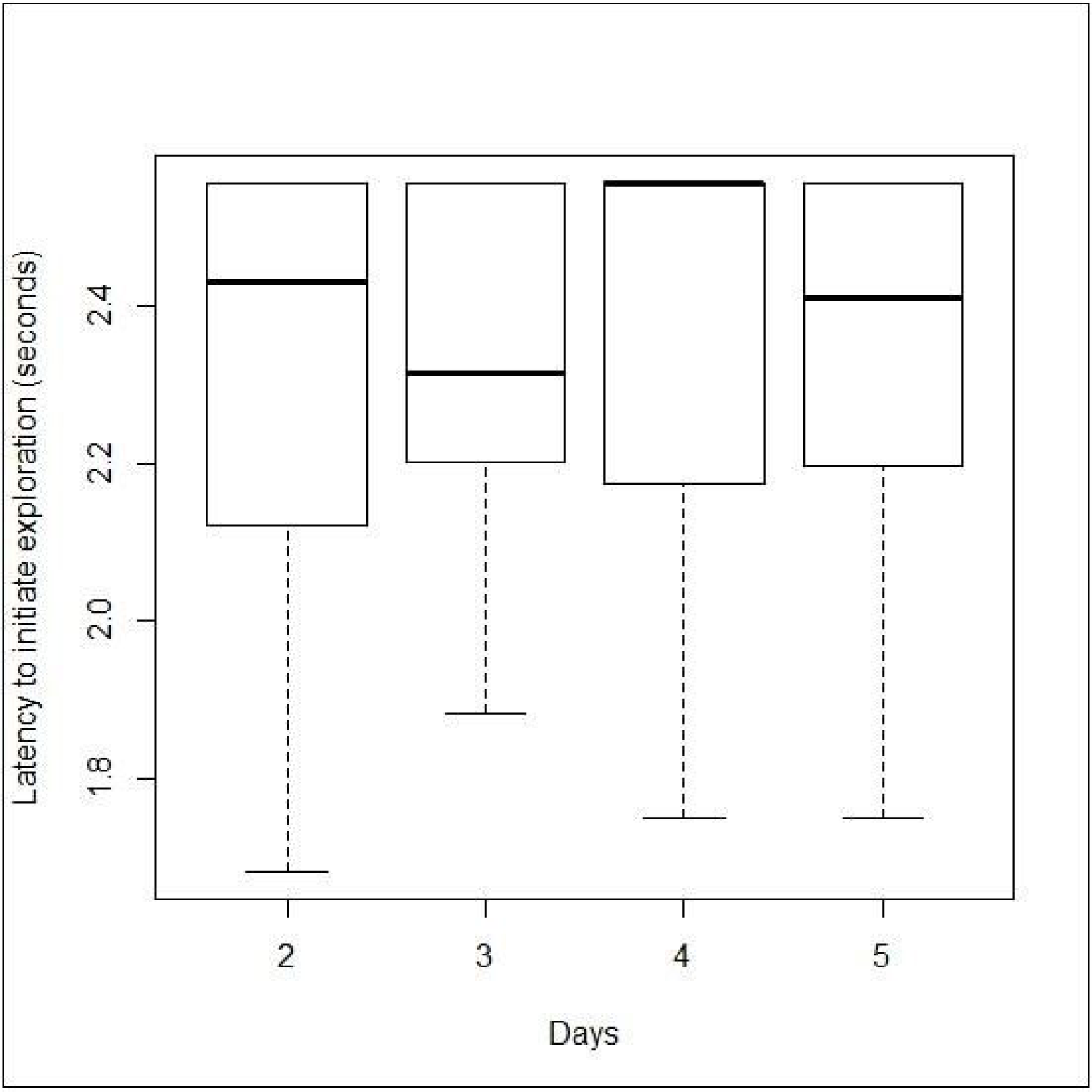
Influence of the repeated exposure for five consecutive days on the boldness of climbing perch in swimway apparatus. The data is log converted and the value for day 1 is represented in Figure 1.

**Figure 3.**
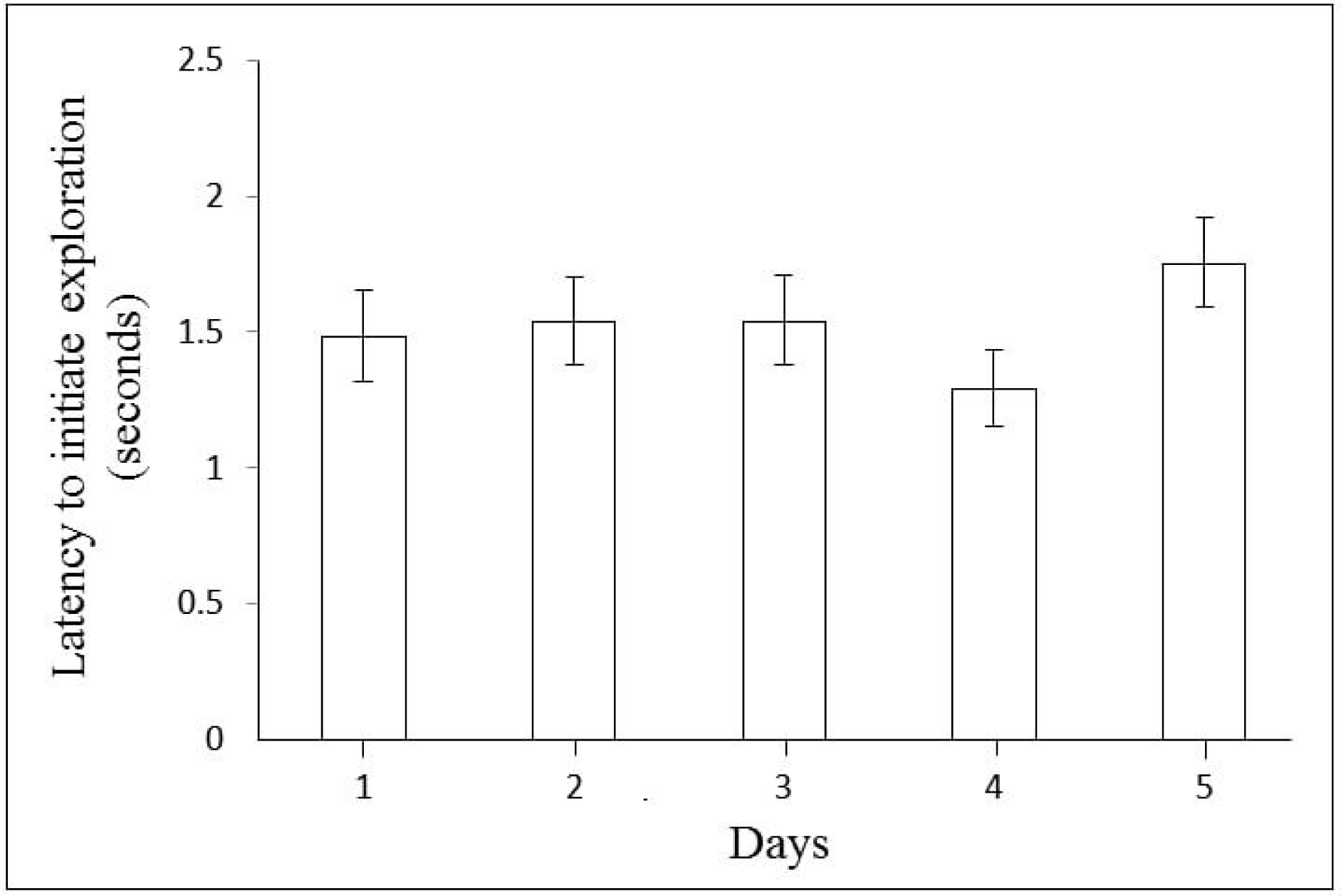
Influence of the repeated exposure for five consecutive days on the boldness of climbing perch in ‘apparatus 1b’. The data is log converted.

**Figure 4.**
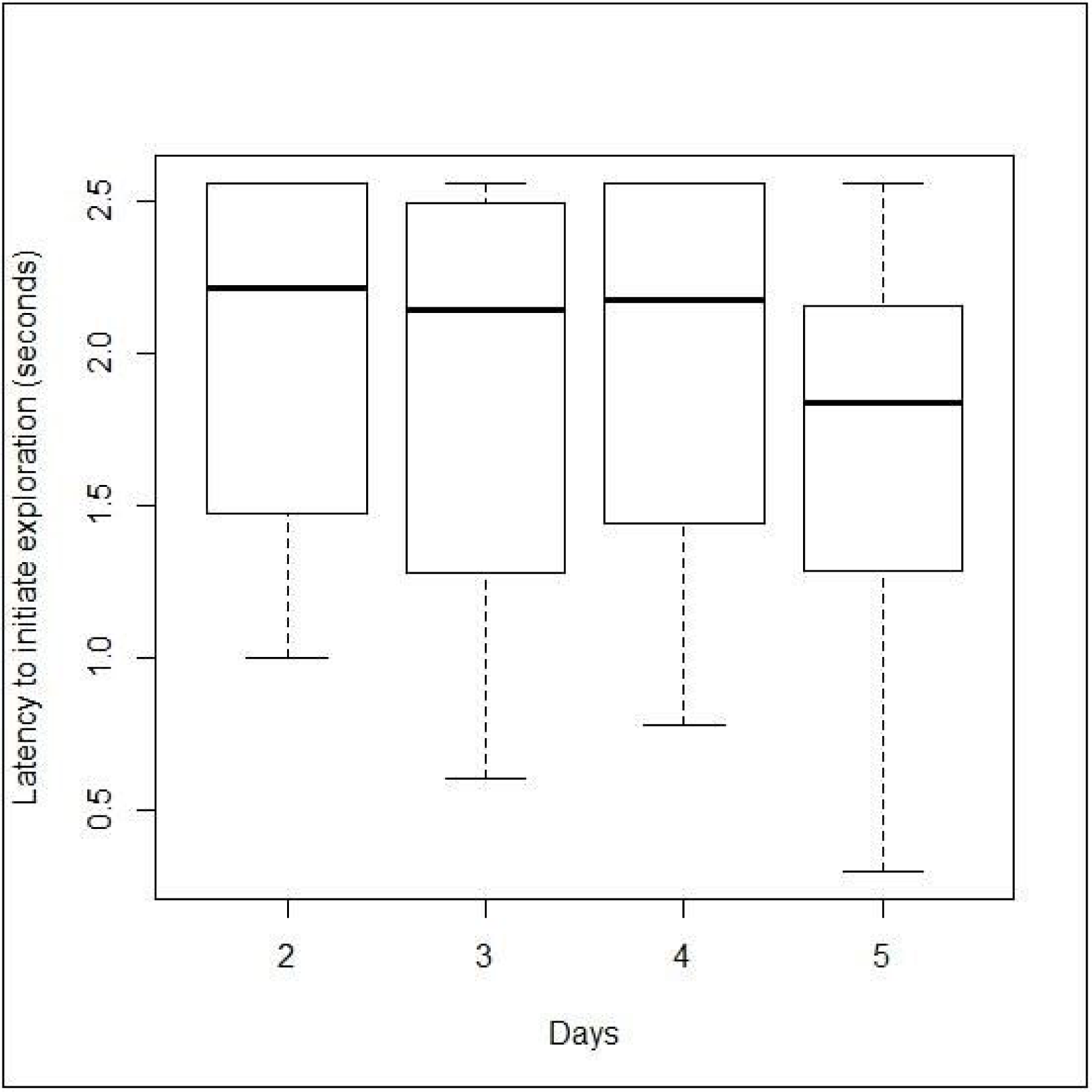
Influence of the repeated exposure for five consecutive days on the boldness of climbing perch in circular open field. The data is log converted and the value for day 1 is represented in Figure 1.

### Experiment 3: Hydrilla cover, cobble substratum and boldness

The modification of start chamber was found to be influencing the boldness of the climbing perch (Repeated measures ANOVA; F_18_ = 3.84, p = 0.01; Fig. 5). The subject fish exhibited significantly reduced interest to leave the start chamber when the floor was covered with cobbles. The presence of the hydrilla cover over the water surface of chamber A did not affect boldness of the subject fish but it neutralised the negative influence of the cobble substratum on the boldness when given in combination with the latter (Table 4).

**Table 4.** Influence of the presence of hydrilla cover and cobble substratum in the start chamber of apparatus 1b on the boldness of the climbing perch - *Post hoc* analysis (Tukey’s test). Statistical significance is denoted as * = p<0.05, ** = p<0.01, *** = p<0.001

|  | **Cobble substratum** | **Hydrilla cover** | **Cobble substratum + Hydrilla cover** |
| --- | --- | --- | --- |
| **Glass substratum** | $t_{18} = 3.62$<br>$p < 0.01^{**}$ | $t_{18} = -0.84$<br>$p > 0.05$ | $t_{18} = 1.55$<br>$p > 0.05$ |
| **Cobble substratum** | — | $t_{18} = -4.46$<br>$p < 0.001^{***}$ | $t_{18} = -2.06$<br>$p > 0.05$ |
| **Hydrilla cover** | — | — | $t_{18} = 2.40$<br>$p > 0.05$ |

**Figure 5.**
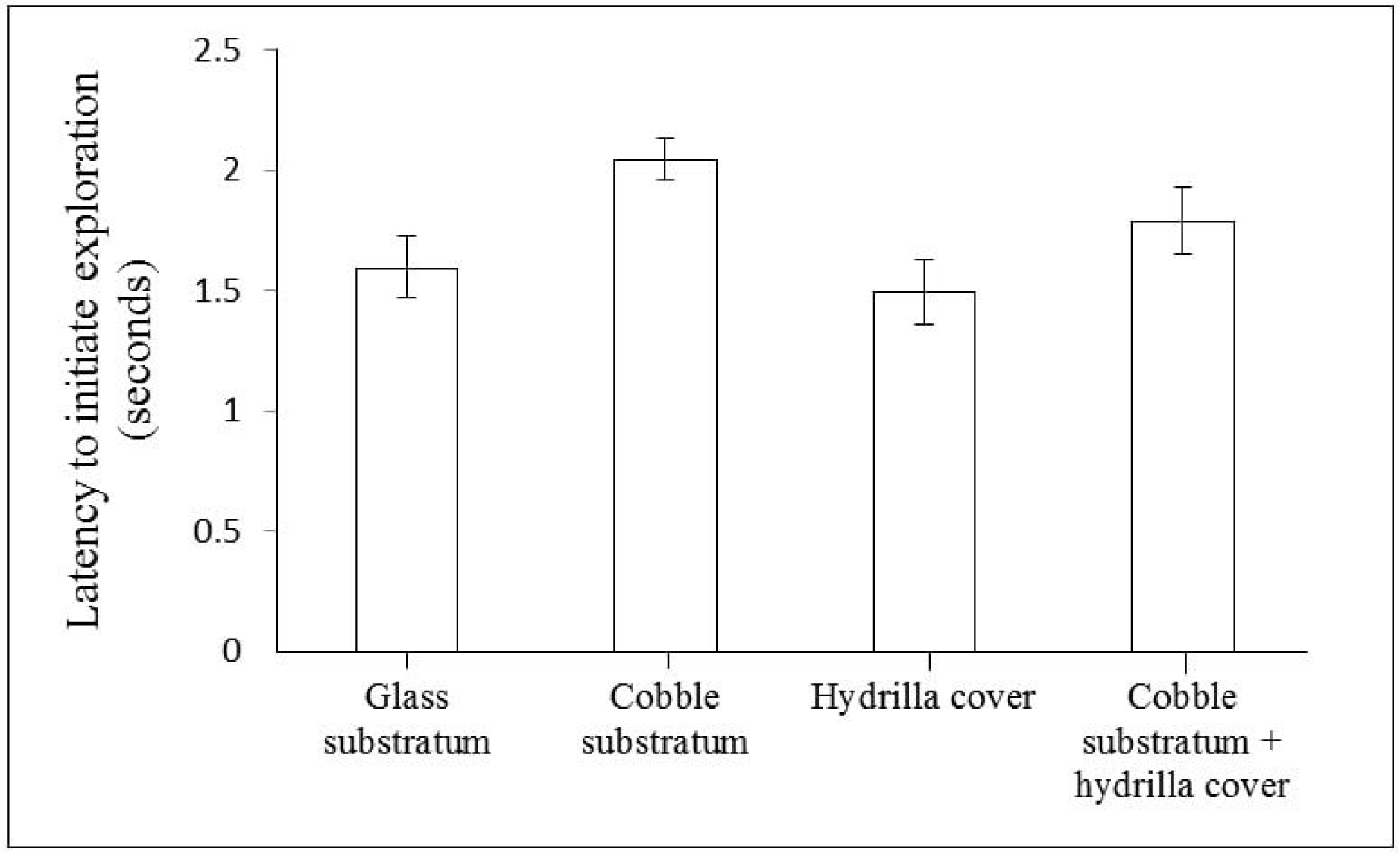
Influence of the presence of hydrilla cover and cobble substratum in the start chamber of apparatus 1b on the boldness of the climbing perch. The data is log converted.

## Discussion

The climbing perch exhibited a noticeable variation in the latency to initiate exploration in different apparatus revealing that boldness in this species is sensitive to the instrument used for quantification. When tested using a swimway, the majority of the subject fish preferred to stay inside the start chamber. However, when the cover present on chamber A was removed (now apparatus 1a), many individuals emerged to explore the well-lit swimway. In apparatus 1a where the start chamber could function as potential shelter (Binoy, 2015), although the shade available inside was much less in comparison to the swimway, the boldness exhibited by the climbing perch was not different from the circular or rectangular open field. This result shows that reduction in the refuge value of the start chamber of the swimway could encourage the subject fish to leave the shelter and extend its activity to the novel areas (Yoshida et al., 2005). The significantly higher level of boldness shown by climbing perch in circular and rectangular open field apparatus, in comparison to the swimway, also points to the impact of the shelter on the boldness in this species. A similar preference for the shaded areas present in the experimental arena, has also been reported in species such as banded killifish (*Fundulus diaphanous*), three spined stickleback (*Gasterosteus aculeatus*), etc. (Dowling and Godin, 2002; Krause et al., 1998). According to Maximino et al., (2010) fishes such as zebrafish (*Danio rerio*), goldfish (*Carassius auratus*), guppy (*Poecilia reticulata*) and tilapia (*Oreochromis niloticus*) exhibit a higher level of scototaxis - the preference for darkness. The question whether scototaxis is a determinant factor of boldness in climbing perch will only be answered by the future studies. Another reason behind the restricted activity in the shelter by the subject fish in the swimway could be the fear generated by the social isolation. Climbing perch being a species preferring to live in groups (Binoy and Thomas, 2004; Binoy, 2015; Zworykin, 2018) separation from the conspecifics may be inducing the antipredator behaviour - staying in the safety of the shelter (Krause et al., 1998; Teplitsky et al,. 2003).

Boldness exhibited by the climbing perch in apparatus 1b - a rectangular open field divided in two unequal chambers using a transparent partition with gate - differed significantly from all other apparatus except apparatus 1a. Here, fish started exploration of chamber B shortly after the introduction into the start chamber. Climbing perch being a species highly dependent on visual cues (Bersa, 1997; Binoy, 2008) it was expected that presence of a transparent partition would not bring a noteworthy change in the boldness expressed in the rectangular open field. Although finding the exact reason behind the increased boldness in the apparatus 1b would require further research, it may be speculated that, as reported in some other species of fishes (Näslund et al., 2015) in climbing perch also the boldness may be sensitive to the changes in the spatial properties of the apparatus. For instance, in wild brown trout (*Salmo trutta*) modification of the size of gate between the start chamber and open area was found to be influencing the tendency of the individuals to extend their activities to the open area (Näslund et al., 2015).

Alteration of the refuge value of chamber A of the apparatus 1b by introducing materials present in the natural habitats of the climbing perch; hydrilla on the water surface and cobbles on the substrata provided an interesting result. When the substratum of the start chamber was changed to cobble, subject fish spent more time in the gaps available between the stone, which reflected as a decrease in the boldness. Here the spatial complexity (Näslund et al., 2015) added by the cobbles to the start chamber may be reducing the boldness of the subject fish by acting as a shelter. Although the presence of the cover on top of the start chamber (swimway) negatively influenced the boldness of the climbing perch, covering the water surface of the start chamber using hydrilla was found to be inefficient in influencing this behavioural trait significantly. From this result it could be assumed that the shade generated by the hydrilla cover in the start chamber might be less in comparison to the swimway since light could penetrate through the gaps between the water plants. Amusingly when hydrilla cover given in combination with the cobble substratum the former cancelled the negative impact induced by the latter on the boldness in this species. This result points to another possible reason for leaving the start chamber with hydrilla cover by the subject fish soon after the introduction. Visiting water surface (Graham, 1997) and gulping atmospheric air is obligatory to the climbing perch, blocking which it may suffer suffocation (Bersa, 1997; Graham, 2006). Hence although the cobbles were providing shelter, climbing perch moved away from the start chamber when the water surface was covered with hydrilla since it could hinder the surfacing and air gulping.

In climbing perch although the latency to initiate exploration came down in all three apparatus tested with the repeated exposure for five consecutive days the pattern of change was not consistent. In the swimway, boldness exhibited on the first day was significantly less in comparison to the other days except the fourth day while in the circular open field such a difference was observed only between the first and fifth day. In this set of experiments too, apparatus 1b stood as an outlier and here the boldness did not change significantly as a result of the repeated exposure. According to Matsunaga and Watanabe (2010) though the relationship between boldness, exploration and habituation had been a topic of intense investigation in rodents, this line of research did not get an equal level of attention from ichthyologists. However, understanding linkage between boldness and habituation is vital since it could give insights into various cognitive phenotypes of the individual such as long term and short term memory (Leussis and Bolivar, 2006; White et al., 2017). The current result points to the need for considering and describing the properties of the apparatus being used for measuring such relationship as well as the caution to be taken while extending results obtained from studies conducted using one kind of apparatus to the others without proper investigation.

## Conclusion

To conclude, results of the present study revealed that the trait boldness in the subject species climbing perch is sensitive to the apparatus being used for the measurement. The swimway apparatus, popularly used for testing the emergence latency in fishes, was found to be less effective in this species since very few individuals came out of the start chamber. Amongst the different apparatus tested, apparatus 1b (rectangular open field divided in two unequal chambers) gave a unique result. The climbing perch exhibited a higher level of boldness in apparatus 1b compared to the other instruments and here the repeated exposure for five days didn’t affect this trait significantly. The potential of the cobble substratum to reduce the boldness and the hydrilla cover to neutralise it, points to the need for designing and using apparatus considering the properties of the natural habitats of subject species for quantifying behavioural and personality traits including boldness in the laboratory. Boldness is a trait with very high impact on the fitness and survival of a fish species but sensitive to various physiological (metabolism, hunger) and ecological parameters such as predation pressure, early life experience, ecosystem variation, etc (Chapman et al., 2010; Harris et al., 2010). Hence future studies focusing on these aspects along with the properties of the experimental arena will elucidate a clear picture of the boldness in climbing perch, a sturdy species surviving in diverse environmental conditions.

## Acknowledgements

VVB is grateful to the Science and Engineering Research Board (SERB), Department of Science and Technology, Government of India, for a grant (SB/FT/LS-155/2012) that enabled this study. The experiments reported in this paper comply with the current relevant laws of India.

